# Place cell firing cannot support navigation without intact septal circuits

**DOI:** 10.1101/470088

**Authors:** Kevin A. Bolding, Janina Ferbinteanu, Steven E. Fox, Robert U. Muller

## Abstract

Though it has been known for over half a century that interference with the normal activity of septohippocampal neurons can abolish hippocampal theta rhythmicity, a definitive answer to the question of its function has remained elusive. To clarify the role of septal circuits and theta in location specific activity of place cells and spatial behavior, three drugs were delivered to the medial septum of rats: tetracaine, a local anesthetic; muscimol, a GABA-A agonist; and gabazine, a GABA-A antagonist. All three drugs disrupted normal oscillatory activity in the hippocampus. However, tetracaine and muscimol both reduced spatial firing and interfered with the rat’s ability to navigate to a hidden goal. After gabazine, location specific firing was preserved in the absence of theta, but rats were unable to accurately locate the hidden goal. These results indicate that theta is unnecessary for location specific firing of hippocampal cells, and that place cell activity cannot support accurate navigation when septal circuits are disrupted.

---

The ability of rats to solve complex spatial problems is thought to depend on a neural navigational system centered on the hippocampus. This conclusion was based on the discovery of “place cells,” which are pyramidal cells of Ammon’s horn (O’Keefe and Dostrovsky, 1971). Since each place cell fires rapidly only when the rat’s head is in a restricted “firing field” and fields are distributed over the environment, the time-averaged, across-cell discharge pattern is unique for each location so that the place cell population acts as a cognitive map. The mechanisms by which the hippocampus updates the place cell representation during locomotion and converts this information to adaptive behavioral output are critically important not only for understanding spatial navigation, but also for the hippocampus’ more general function in memory-guided behavior (Scoville and Milner, 1957; O’Keefe and Nadel, 1978).

The place cell firing rate vector must be continuously modified during locomotion, and this computational demand coincides with the appearance of a distinct hippocampal activity state dominated by the 5 – 12 Hz theta rhythm (Vanderwolf, 1969). On average, place cell discharge is phase-locked to theta (Fox et al., 1986), and place field firing is more rapid during theta than otherwise (Kubie, 1985). Place cells also show “phase precession” in which spikes occur at progressively earlier theta phases as the rat proceeds through the firing field (O’Keefe and Recce, 1993). Models have demonstrated how theta oscillators could contribute directly to the formation and expression of place cell firing fields (Lengyel et al., 2003; Welday et al., 2011). Alternatively, theta may have little direct role in producing spatially-localized firing (Brandon et al., 2011; Yartsev et al., 2011; Brandon et al., 2014), but may allow the hippocampus to encode spatial or temporal event sequences (Gupta et al., 2012; Wang et al., 2015) or to efficiently broadcast spatial information to downstream structures (Tabuchi et al., 2000; Benchenane et al., 2010; van der Meer and Redish, 2011; Tingley and Buzsaki, 2018). Therefore, to assess the functional relevance of hippocampal theta, it is of great interest to attenuate or abolish the rhythm independent from effects on place cell activity.

Hippocampal theta depends on a ‘pacemaker’ circuit formed by reciprocal inhibitory connections within and between the medial septum and the hippocampus (Freund and Antal, 1988; Toth et al., 1993; Wang, 2002). Temporary or permanent lesions of the medial septum disrupt hippocampal theta (Allen and Crawford, 1984; Smythe et al., 1992; Bland et al., 1996; Koenig et al., 2011) and impair performance in hippocampus-dependent tasks (Winson, 1978; Mizumori et al., 1989; Givens and Olton, 1990; Mizumori et al., 1990; Givens and Olton, 1994), but it remains unclear whether the behavioral deficits are caused directly, by loss of theta rhythmicity *per se*, or indirectly, through degraded place cell activity (Mizumori et al., 1989).

Understanding the functional role of theta oscillations requires manipulations of theta rhythm that do not affect overall hippocampal activity. To address this issue, we compared septal injections of three drugs with distinct mechanisms of action in their ability to reduce theta, the concomitant effects on place cells and their disruption of a spatial task. Two of the drugs suppress neural activity, producing reversible lesions of the septal circuit: tetracaine, acting as a local anesthetic, blocks action potentials, and muscimol a GABA-A agonist, inhibits local circuit activity. The third drug, the GABA-A receptor antagonist gabazine, removes inhibition within the septal circuit. Here we demonstrate that infusing gabazine into the medial septum produces robust theta blockade, while leaving CA1 and CA3 place cell activity intact, in contrast to effects of septal inactivation. Despite its selective effect on theta rhythm, gabazine produced the same spatial memory deficit as tetracaine and muscimol. Therefore, hippocampal place cell activity is not sufficient for navigation. Instead, the ability to convert place cell firing into adaptive spatial behavior depend on intact circuits within the medial septum and correlates with the strength of hippocampal theta rhythm.

## Results

### Hippocampal theta oscillations are reversibly suppressed by septal injections of tetracaine, muscimol or gabazine

To determine how hippocampal activity depends on the integrity of medial septal circuits, we acquired tetrode recordings from the hippocampus of foraging rats before and after septal injections of tetracaine (local anesthetic), muscimol (GABA-A agonist), gabazine (GABA-A antagonist), or the vehicle (PBS) used to dissolve the three active substances (Figure 1). To identify the maximal effects of each drug on hippocampal theta, we found the period of minimum theta power in a sliding 16-min time window after drug infusion for each recording session. To exclude large irregular LFP activity associated with immobility, we included only data recorded while the animal was moving at a speed greater than 3 cm/s in this analysis. Because searching for minima inherently yields lower than average values, we compared these values to minima computed for each baseline session using an 8-min sliding window. Minimum theta power in baseline sessions was 92.42 ± 7.37% (mean ± s.d., n = 30 sessions) of overall baseline theta power. Hippocampal theta power was moderately reduced after septal PBS infusions (Figure 1A and E, minimum theta: 77.89 ± 7.88%, mean ± s.d., n = 9 sessions). For 7/9 PBS injections, the period of greatest theta reduction was in the first 32-minute post-injection session (Figure 1E). Thus, volumetric injections may temporarily alter normal activity in medial septal circuits and subtly reduce theta entrainment of the hippocampal LFP.

**Figure 1.**
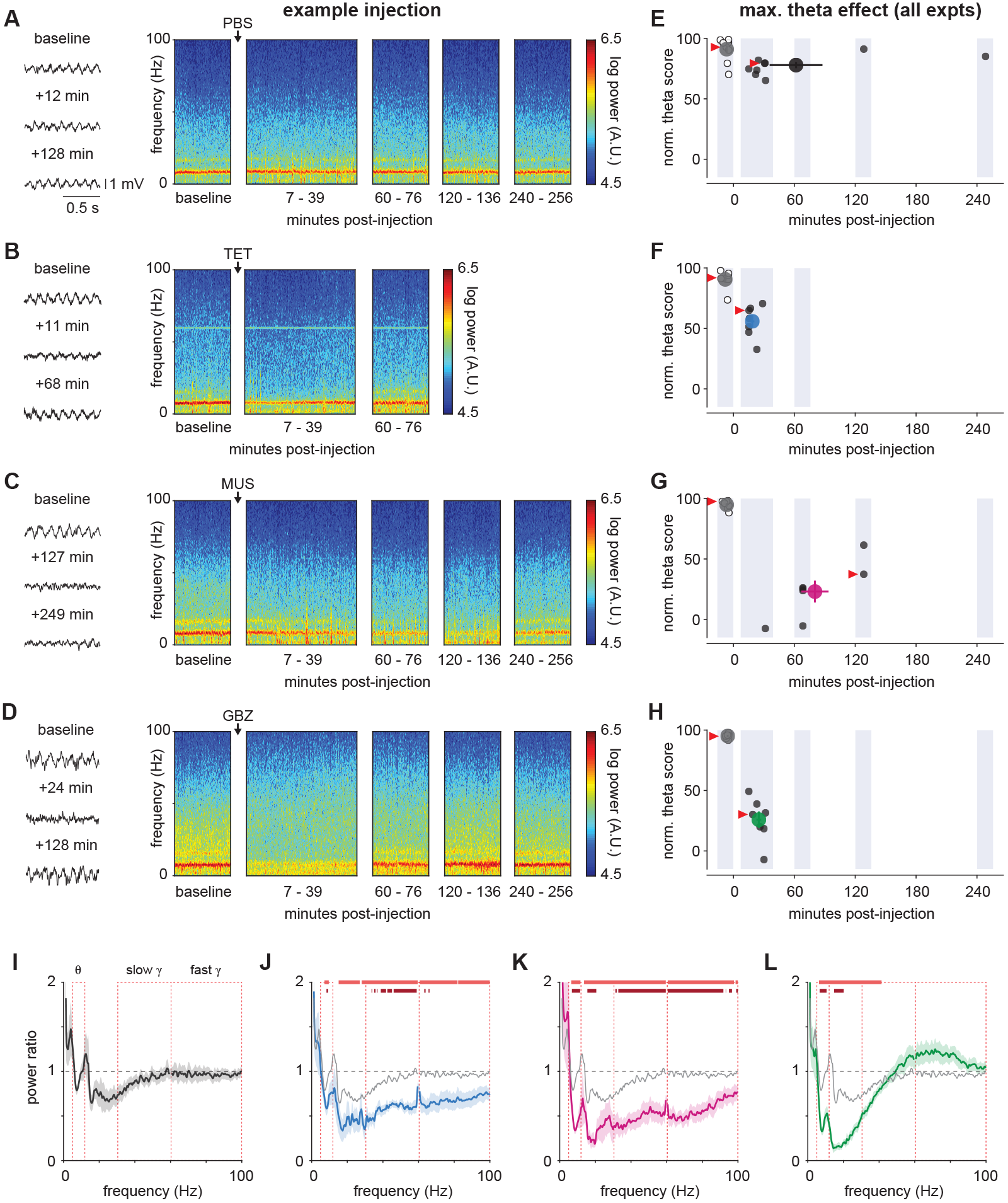
Septal inactivation or GABA-A antagonism degrades hippocampal theta oscillations. (A-D) Changes in hippocampal LFP after septal injections of PBS (A), tetracaine (B), muscimol (C), or gabazine (D). Left, Examples of LFP traces recorded during baseline sessions and post-infusion recording sessions for each drug. The time when the LFP trace was recorded is indicated. Right, Spectrograms for the baseline and subsequent sessions for the corresponding injection experiment. Post-infusion effects were evaluated starting at 4 different points in time: t = 7 min [32-min recording], t = 60 min [16-min], t = 120 min [16-min], and t = 240 min [16-min]). For tetracaine, sessions starting at t = 120 and t = 240 min were omitted because preliminary work indicated that its effects were no longer detectable at the end of the first 32-min session. (E-H) Summary of time course and depth of maximal theta reduction following septal injections of PBS (E, n = 9, paired t-test, post-injection vs. baseline minimum: p = 0.02), tetracaine (F, n = 7, p = 4.02e-5), muscimol (G, n = 7, p = 1.14e-4), or gabazine (H, n = 7, p = 0.001). Empty circles represent the normalized theta score at the middle of the 8-minute interval during maximal theta reduction for each experiment in the baseline session. Filled circles are the normalized theta score at the middle of the 16-minute interval during maximal theta reduction post-injection. Large, filled circles and error bars show the average score and timing of minimal theta (mean ± s.e.m). Red arrowheads mark the sessions used to generate the example spectrograms in A-D. (I-L) Average power ratios between post-injection and baseline power spectra (mean ± s.e.m). (I) Power spectrum was not significantly affected by PBS infusion (no significant difference from 1 for any frequency). (J-L) Light red line indicates p < .05, t-test vs. ratio of 1. Dark red line indicates p < .05, unpaired t-test vs. power ratio for PBS. P-values are corrected for multiple comparisons using the false discovery rate. To facilitate direct visual comparison, power ratios for PBS are overlaid in gray.

To suppress medial septal input to the hippocampus, we used two substances with distinct mechanisms of action: tetracaine, a local anesthetic; and muscimol, a GABA-A receptor agonist. Septal tetracaine injections produced a rapid degradation in hippocampal theta power, often apparent as soon as recording resumed at 7-min post-injection (time to max effect: 18.6 ± 5.3 min. post-injection; mean ± s.d., n = 7 sessions). Maximum theta reduction after tetracaine injection was significantly greater than after PBS (Figure 1B and F; minimum theta: 55.89 ± 13.29%, mean ± s.d., n = 7 sessions, p = 9.91e-04, unpaired t-test vs. PBS). Tetracaine effects also had rapid offset kinetics; theta power normally recovered by the start of the 16-min recording session at t = 60-min post-infusion. In contrast to the rapid tetracaine action, the reduction of theta following muscimol infusions had a slow onset and delayed recovery (time to max effect: 80.3 ± 35.6 min. post-injection; mean ± s.d., n = 7 sessions). For 6/7 muscimol injections the maximal effect occurred 1 hour or more post-infusion, and recovery was only partial at the end of the last recording session 4 hours post-infusion. The maximum theta reduction was greater than for tetracaine (Figure 1C and G; minimum theta: 23.04 ± 23.94%, mean ± s.d., n = 7 sessions). No systematic changes in running speed accompanied the theta power reductions induced by tetracaine and muscimol injection (Figure 2).

**Figure 2.**
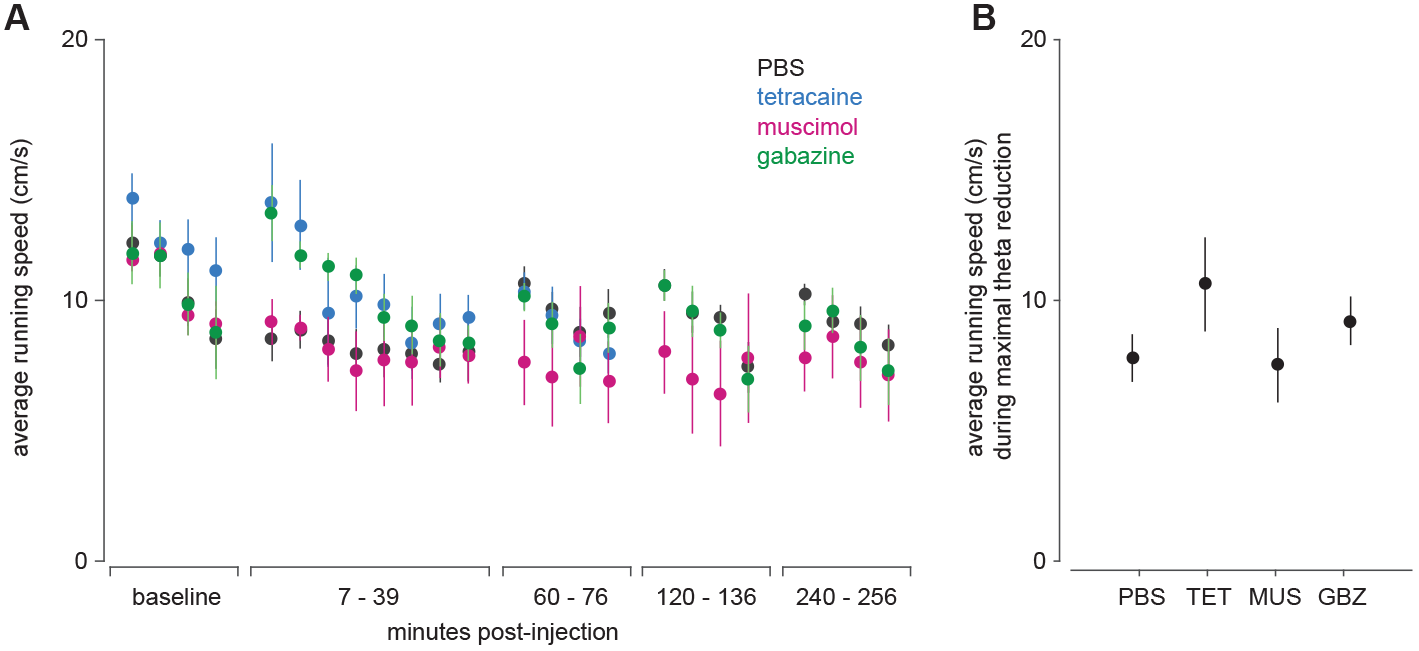
Running speeds were not systematically reduced during maximal theta reduction. (A) Average running speeds in non-overlapping 4-min windows during baseline and post-injection sessions for each drug (mean ± s.e.m). (B) Average running speeds during the 16-min period of maximal theta reduction used in LFP and place cell analysis. (ANOVA for effect of drug: F(3,26) = 1.22, p = 0.32). Note that tetracaine and gabazine tended to increase running speeds during the period when hippocampal theta power was maximally reduced; however, these effects did not result in statistically significant differences.

Tetracaine and muscimol were effective in disrupting hippocampal theta oscillations but may have additional effects due to the silencing of excitatory and neuromodulatory septohippocampal afferents. To identify how hippocampal function depends specifically on theta rhythm we sought a means to disrupt theta using a distinct circuit mechanism. Infusion of a GABA-A receptor antagonist into medial septum increases the activity (Brazhnik, 2004) and reduces the rhythmicity (Brazhnik and Fox, 1999) of both short spike (putative GABAergic) and long spike (putative cholinergic) septohippocampal neurons. Therefore, we also recorded in hippocampus during and after medial septal injections of gabazine, a GABA-A receptor antagonist. Gabazine injections produced large and rapid-onset degradation of theta power in the hippocampal LFP (Figure 1D and H; minimum theta: 25.87 ± 18.01%, mean ± s.d., n = 7 sessions). In general, gabazine caused a longer lasting decrease of hippocampal theta power than tetracaine, but it had a briefer effect and more rapid onset than muscimol, with no associated change in running speed (time to max effect: 25.0 ± 6.2 min. post-injection; mean ± s.d., n = 7 sessions). Thus, blockade of GABA receptors in medial septum was as effective in suppressing hippocampal theta power and more temporally-restricted than septal inhibition through GABA-A receptor activation.

Septal inactivation or disinhibition may broadly affect hippocampal oscillations even outside the theta range through changes in the multiple neurotransmitter systems comprising the septohippocampal projection. Therefore, we examined power ratios before and after drug injection across the frequency spectrum. Each ratio was obtained by dividing the average power during the 16-min period of maximum theta decrease by the average power during the baseline 16-min session. PBS infusions only subtly affected hippocampal oscillations (Figure 1I). In contrast, both muscimol and tetracaine reduced power ratios across the frequency spectrum compared to baseline, including the slow (30-60 Hz) and fast (60-100 Hz) gamma ranges (Figure 1J and K). Gabazine also significantly reduced slow gamma power in a restricted range below 40 Hz, and there was a nonsignificant trend toward a positive effect on fast gamma power (Figure 1L). These data indicate that septal inactivation broadly inhibits oscillatory activity in the hippocampus, whereas septal disinhibition disrupts oscillations in a more specific (5-40 Hz) range.

### The effects of septal inactivation vs. septal disinhibition on place cell discharge

We next asked how septal injections affect the activity and spatial coding of hippocampal place cells. Place cell activity during 16 min baseline sessions was compared to activity during the 16 min intervals for which theta power was maximally reduced by septal injections. Data were included only when the rat explored > 80% of the cylinder area during both sessions. To be analyzed, a cell had to be classified as a place cell during the baseline session.

The majority of place cells displayed theta-modulated rhythmic firing patterns in the baseline session (CA1: 77.1% (101/131 cells); CA3: 50.1% (56/111 cells); theta mod. score > 5; see Methods; Figure 3A-D). We examined drug-induced changes in theta-modulation using the ratio of post-injection to baseline theta-modulation scores. Septal inactivation (tetracaine and muscimol) or disinhibition (gabazine) reduced theta modulation in both CA1 and CA3 cells by >50%, with CA3 cells relatively more affected than CA1 cells after septal inactivation (Figure 3E). Interestingly, tetracaine produced much larger changes in place cell theta modulation than it did on the theta oscillations in the local field potential (Figure 1F), suggesting the remaining rhythmic input is not sufficient to support cell-autonomous theta oscillations.

**Figure 3.**
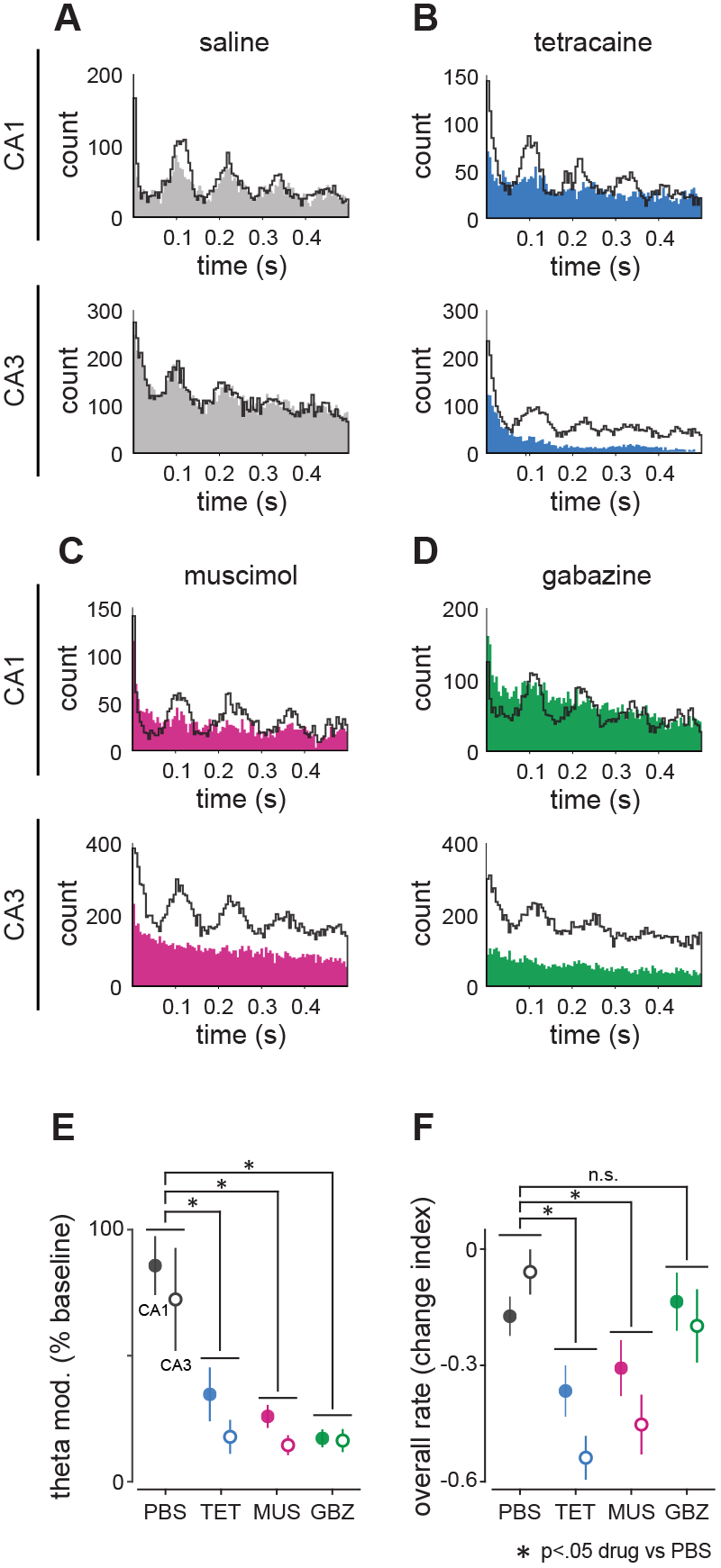
Septal injections disrupt rhythmic discharge of hippocampal place cells. (A-D) Autocorrelograms for CA1 and CA3 place cells before (black line) and after (filled area) intraseptal infusions of each substance. Saline infusions had little effect on the amplitude of successive peaks. In contrast, theta discharge rhythmicity was drastically reduced, and in some cases abolished, by injections of tetracaine, muscimol or gabazine. (E) Change in average theta modulation score for CA1 and CA3 place cells caused by septal injections (mean ± s.e.m.). 2-way ANOVA: [drug: F(3, 149) = 13.94, p = 4.64e-8; region: F(1, 149) = 5.44, p = 0.02; drug X region: F(3, 149) = 0.33, p = 0.8]. Sample sizes (n cells): PBS, CA1: 11, CA3: 26; tetracaine, CA1: 19, CA3: 17; muscimol, CA1: 30, CA3: 14; gabazine, CA1: 26, CA3: 14. Circles are mean ± s.e.m. Filled circles: CA1; empty circles: CA3. Asterisks indicate p < .05 for post-hoc unpaired t-tests. No within-drug region effects were significant. (F) Change index [post - baseline / post + baseline] for overall place cell firing rates between baseline session and post-injection during maximal theta reduction (mean ± s.e.m.). 2-way ANOVA: [drug: F(3, 234) = 10.28, p = 2.21e-6; region: F(1, 234) = 1.71, p = 0.19; drug X region: F(3, 234) = 1.53, p = 0.21]. Sample sizes (n cells): PBS, CA1: 34, CA3: 20; tetracaine, CA1: 27, CA3: 41; muscimol, CA1: 37, CA3: 25; gabazine, CA1: 33, CA3: 25.

Modulation score changes might result from reduced activity rather than loss of rhythmic modulation. Strikingly, tetracaine and muscimol produced substantial reductions in firing rate, but the more powerful suppression of theta by gabazine did not substantially alter place cell firing rate (Figure 3F). Nevertheless, drug-induced changes in theta-modulation remained robust when we restricted theta-modulation analysis to only units whose post-drug firing rate was at least 90% of their baseline rate (Figure 4). We conclude that each drug greatly reduced theta frequency modulation of individual cells’ activity when LFP theta was reduced, while septal inactivation had additional suppressive effects on overall firing rates in the hippocampus.

**Figure 4.**
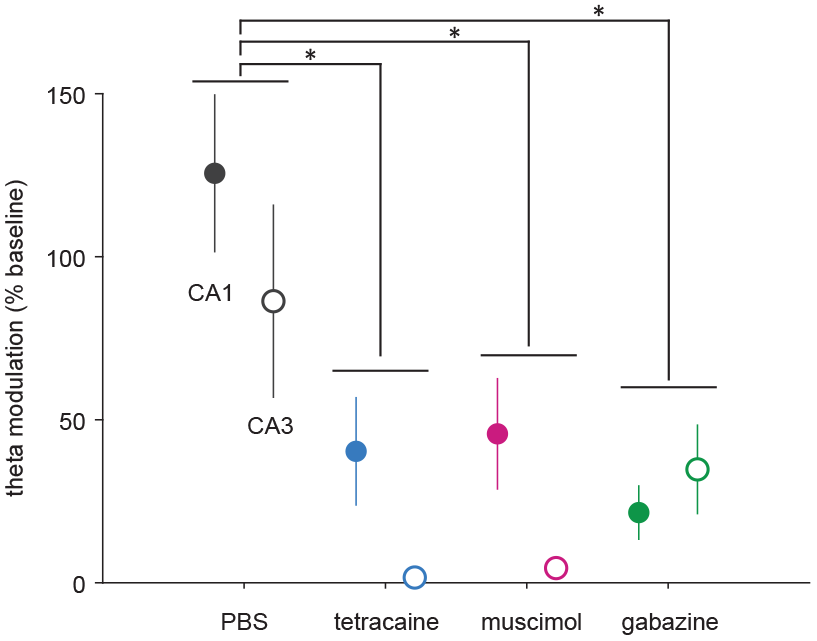
Theta modulation with preserved firing rates. Change in average theta modulation score for CA1 and CA3 place cells caused by septal injections (mean ± s.e.m.) for cells with firing rates ≥90% of baseline. Sample sizes (n cells): PBS, CA1: 11, CA3: 10; tetracaine, CA1: 12, CA3: 5; muscimol, CA1: 7, CA3: 5; gabazine, CA1: 17, CA3: 8. 2-way ANOVA: [drug: F(3, 41) = 8.16, p = 2.24e-4; region: F(1,41) = 2.55, p = 0.12; drug X region: F(3, 41) = 0.71, p = 0.55]. Asterisks indicate p < .05 for post-hoc unpaired t-tests. No within-drug region effects were significant.

We next visualized the effects of intraseptal injections on location-specific firing by comparing firing rate maps from baseline and post-injection recordings (Figure 5). CA1 and CA3 place cells continued to fire at similar rates and in similar locations after vehicle injection (Figure 5A). If theta rhythmic oscillations in individual cells or in the local network are necessary for expressing spatial firing fields, then each of the three drugs should uniformly disrupt place cell firing. Conversely, if spatial firing fields depend on septal output more generally, and do not require specific theta rhythmic entrainment, effects of gabazine may differ from those of tetracaine and muscimol.

**Figure 5.**
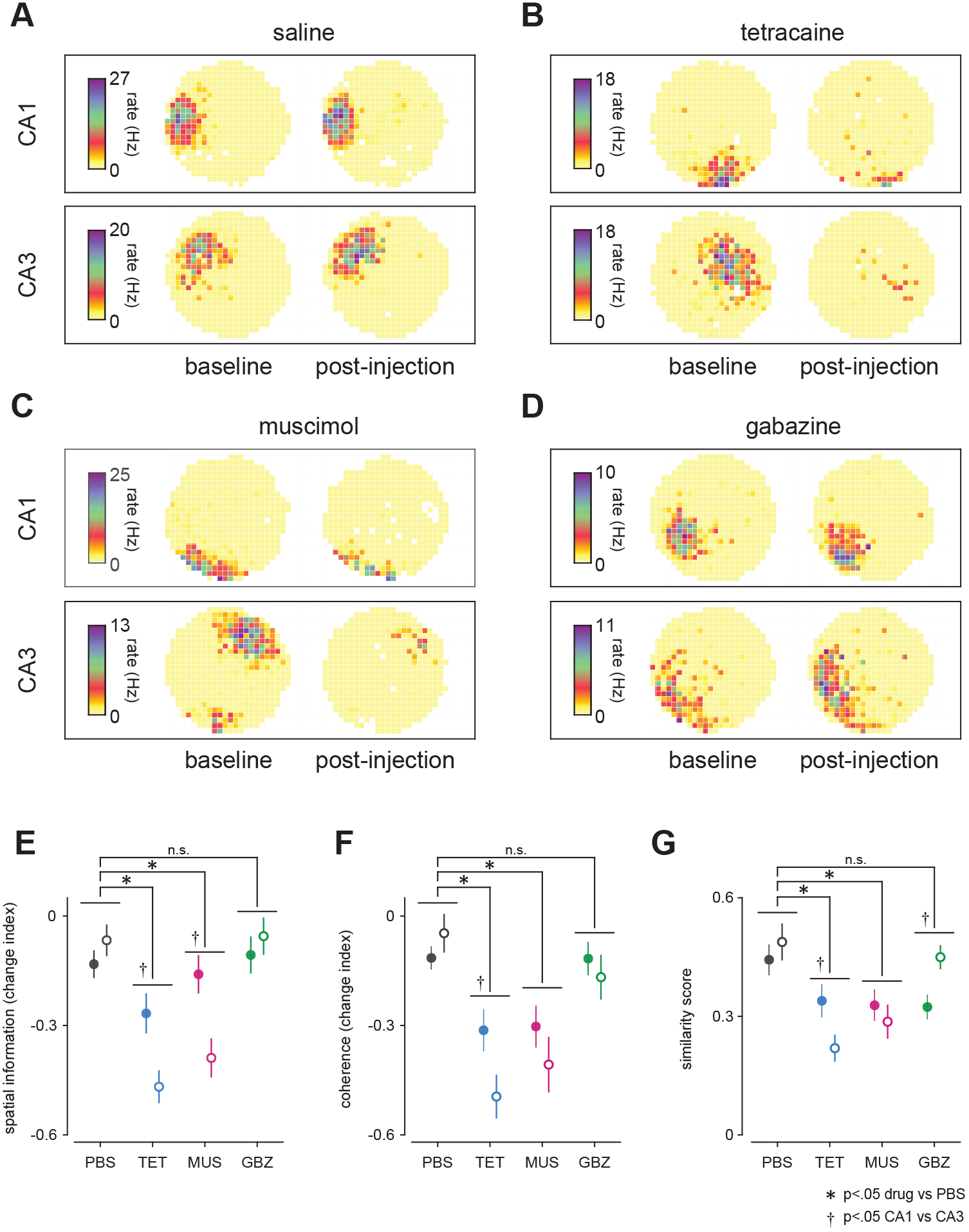
Degraded place cell firing fields after septal inactivation but not septal disinhibition. (A-D) Firing rate maps for one CA1 and one CA3 cell are shown for each substance. Rate reduction was clearly visible after septal injections of tetracaine (B) and muscimol (C) with residual firing at the location of the pre-injection field. Gabazine infusions (D) did not produce visible effects in either CA1 or CA3. (E) Change index [post - baseline / post + baseline] for mean spatial information between baseline session and post-injection during maximal theta reduction (mean ± s.e.m.). 2-way ANOVA: [drug: F(3, 234) = 15.13, p = 4.95e-9; region: F(1, 234) = 4.68, p = 0.03; drug X region: F(3, 234) = 4.78, p = 0.003]. Sample sizes (n cells): PBS, CA1: 34, CA3: 20; tetracaine, CA1: 27, CA3: 41; muscimol, CA1: 37, CA3: 25; gabazine, CA1: 33, CA3: 25. (F) Same as (E) for place cell firing field coherence. 2-way ANOVA: [drug: F(3, 234) = 14.87, p = 6.83e-9; region: F(1, 234) = 2.89, p = 0.09; drug X region: F(3, 234) = 1.71, p = 0.17]. (G) Similarity of spatial firing patterns between baseline session and during maximal theta reduction. 2-way ANOVA: [drug: F(3, 234) = 8.87, p = 1.36e-5; region: [F(1, 234) = 1.51e-3, p = 0.97; drug X region: F(3, 234) = 3.90, p = 0.01]. Filled circles: CA1; empty circles: CA3. Asterisks indicate p < .05 for post-hoc unpaired t-tests. Daggers indicate p < .05 for within-drug region effects.

Indeed, tetracaine and muscimol produced substantial degradation of spatial firing in CA1 and CA3 cells, visible as both a decrease in the peak rate and in the field area (Figure 5B and C). These effects were most consistent following tetracaine and most severe in the CA3 region. In contrast, place cell firing fields were largely intact following gabazine injection (Figure 5D). This pattern of post-injection firing rate changes was also observed in a small population of simultaneously recorded inter-neurons (Figure 6). Consistent with previous reports, firing field locations were, in general, not perturbed by injections of any of the active drugs; even if the firing rate was drastically reduced, residual discharge was at the location of the baseline field (see Figure 5B in CA1).

**Figure 6.**
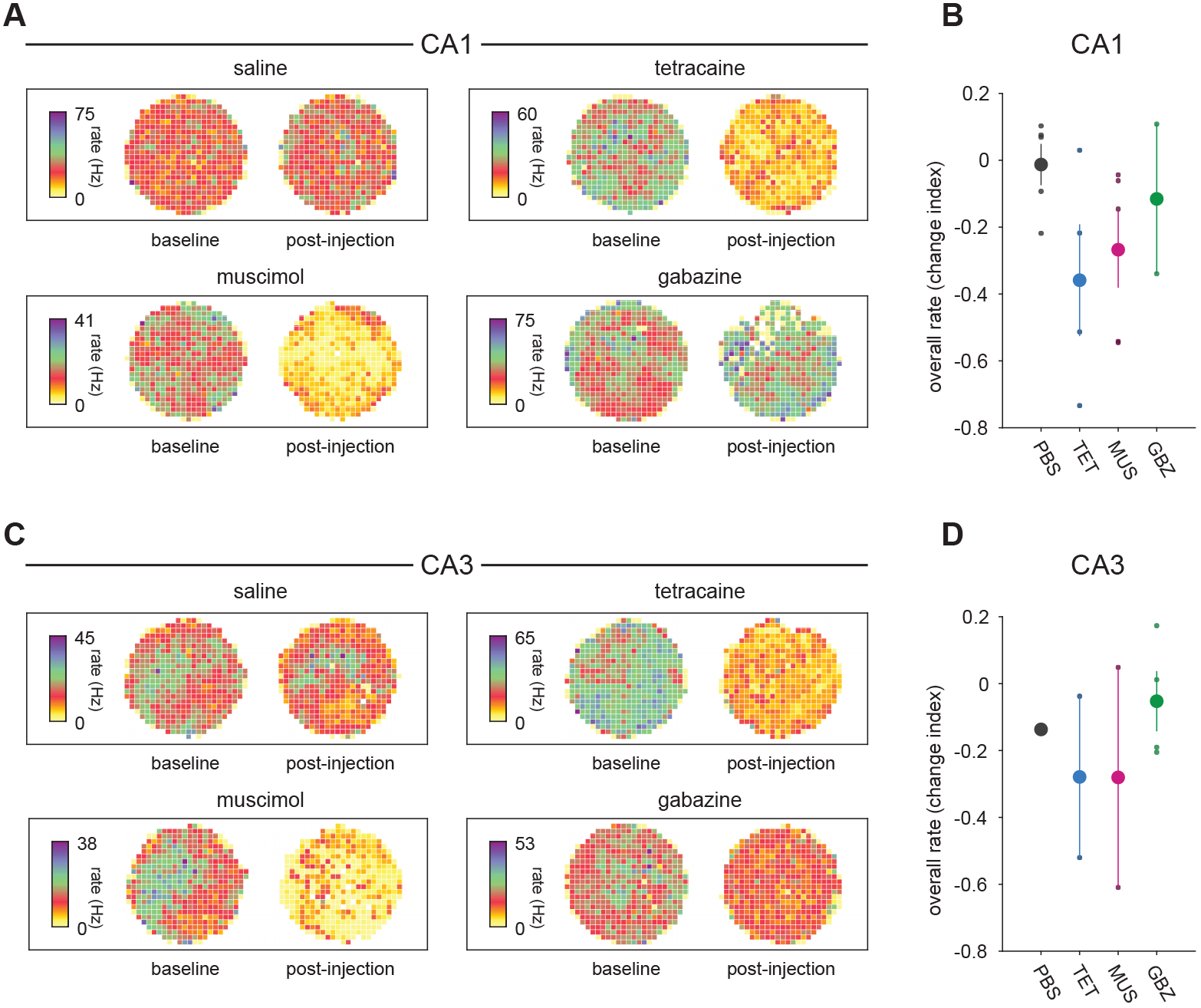
Reduced firing rates in hippocampal interneurons during septal inactivation. (A) Firing rate maps for interneurons recorded in CA1 (overall rate > 4 Hz in baseline session) before and after injection of each substance. (B) Change index [post - baseline / post + baseline] for overall interneuron firing rates between baseline session and post-injection during maximal theta reduction. Large circles are mean ± s.e.m. Small circles show values for individual interneurons. (C-D) As in A-B, but for interneurons recorded in CA3. Sample sizes (n cells): PBS, CA1: 5, CA3: 1; tetracaine, CA1: 4, CA3: 2; muscimol, CA1: 5, CA3: 2; gabazine, CA1: 2, CA3: 4.

To identify reliable changes in spatial firing quality after septal injections, we calculated average spatial information (see Methods; Olypher et al., 2003) and coherence, a measure of the smoothness of place cell firing fields, across the population of CA1 and CA3 place cells and compared changes in these measures to changes after PBS injection. Septal inactivation, either using tetracaine or muscimol, caused significant reductions in both spatial information and coherence, and these effects were consistently stronger in CA3 than CA1 (Figure 5E and F). By contrast, gabazine injections had no significant effect on either spatial information or coherence. Degraded place cell activity after septal inactivation affected the reliability of location-specific firing. Thus, the similarity between baseline and post-injection firing rate maps was significantly lower after tetracaine or muscimol than after PBS, and CA3 cells were affected more strongly than CA1 cells (Figure 5G). Gabazine injections, on the other hand, had no effect on firing field similarity scores.

Overall, the pattern of results is consistent with the conclusion that gabazine leaves place cell firing patterns intact, in contrast to muscimol or tetracaine. It is particularly striking that tetracaine, which produces relatively weak and short-lived effects on LFP theta oscillations, severely disrupted place cell activity, while gabazine, which more completely reduces LFP theta had no significant effects on the measured aspects of place cell coding. These contrasting effects reveal that the strength of the hippocampal theta oscillation can be dissociated from the expression of location-specific firing.

### Effects of septal inactivation vs. septal disinhibition on spatial navigation

The preserved place cell activity after gabazine injections demonstrates that the neural computation of self-localization information occurs independent of the strength of the hippocampal theta rhythm. However, the theta rhythmic organization of hippocampal output may be critical for its ability to communicate to downstream structure and influence ongoing behavior (Tabuchi et al., 2000; Hyman et al., 2005; Benchenane et al., 2010; Tingley and Buzsaki, 2018). To directly assess navigation abilities following septal injections, we trained rats in a place accuracy task where the rats were required to pause in a specific goal area in the recording arena to trigger the release of a sugar pellet from an overhead pellet feeder. Performance was assessed in visible and hidden goal versions of the task. In the visible goal version, the goal area was marked by a disk placed on the floor of the arena. In the hidden goal version, the unmarked goal area had to be found by navigation relative to distal cues (the polarizing cards), presumably using hippocampal cognitive mapping computations. The nature of the task and conclusions drawn by comparing performance in the two versions strongly resemble the water maze paradigm (Morris, 1981; McDonald and White, 1994; Packard et al., 1994). In our task, trained animals repeatedly navigated to the target location resulting in a large peak in the occupancy map in both task versions (Figure 7A & D). Accuracy, measured as time spent in the target region, was nearly equal in baseline sessions across the two versions of the task.

**Figure 7.**
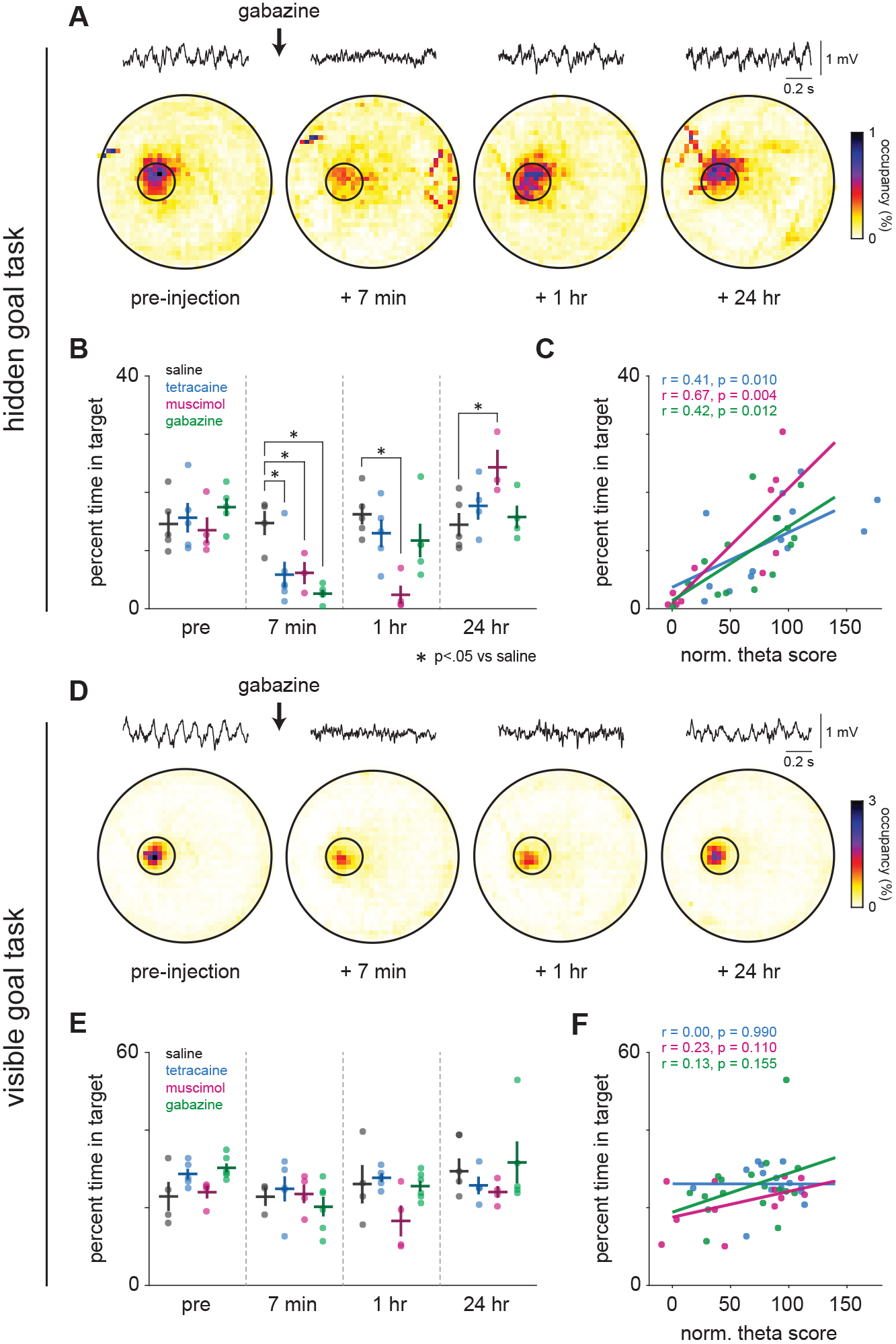
Accurate spatial navigation degraded with either septal inhibition or septal disinhibition. (A) Example of behavioral performance in the hidden goal place-accuracy task during an individual baseline session and the 3 corresponding post-injection sessions. Heatmaps represent percent time spent in each location across a 20-minute session. Small circles mark the hidden target area; large circles represent the arena. Examples of LFP traces from each session are shown above. The time when each of the post-injection sessions began relative to the end of the gabazine injection is indicated below. (B) Time course of spatial accuracy performance before and after injection of PBS (n = 5 experiments), tetracaine (n = 5), muscimol (n = 4), or gabazine (n = 4). Circles represent data from individual experiments. Bold lines are mean ± s.e.m. (C) Correlation of behavioral performance in the hidden goal task and theta power for each substance. Each point represents a normalized theta score and the corresponding level of behavioral performance for one session of one experiment. Lines indicate the linear fit through all points for each substance; recovery of theta oscillations was associated with significantly improved behavioral performance in all cases. (D-F) Same as A-C for the visible goal version of place-accuracy task. (PBS, n = 4; tetracaine, n = 5; muscimol, n = 4; gabazine, n = 6). None of the drugs affected the animals’ ability to perform accurately; and there was no significant relationship between the normalized theta score and time spent in the goal area.

Animals were trained to criterion in both versions of the task (first in the visible goal version and then in the invisible goal version) and then received a series of septal injections (vehicle, gabazine, muscimol, and tetracaine), first in the hidden goal version, followed by the same injection series in the visible goal version; injections were separated by two days. Each experiment consisted of four 20 min sessions: baseline, t = 7 min post-injection (when gabazine and tetracaine have their maximal LFP effect), t = 60 min post-injection (allowing time for muscimol effects to reach their peak) and twenty-four-hour post-injection recovery. In a subset of experiments, animals exhibited either excessive thigmotaxis or immobility after injection; the data from these sessions were not included in further analyses (Figure 8).

**Figure 8.**
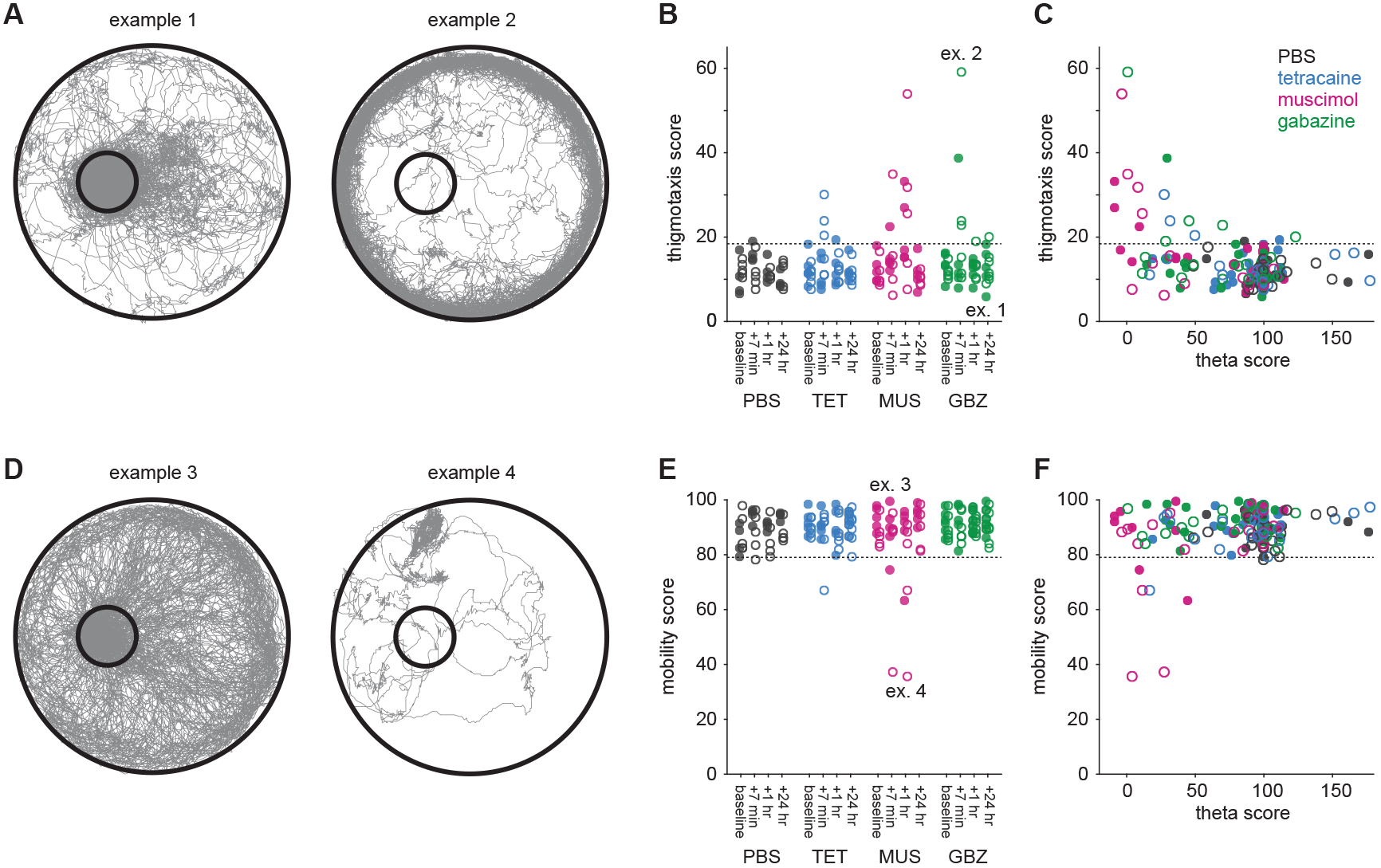
Exclusion criteria for non-specific behavioral effects in place-accuracy sessions. In some place-accuracy task sessions, animals exhibited either pronounced thigmotaxis or immobility after drug injection. (A) Gray lines represent animal trajectories over a 20-minute session, showing sessions with minimum (left, example 1) or maximum (right, example 2) thigmotaxis scores. (B) Thigmotaxis scores across all drugs and sessions. Empty circles show hidden goal sessions, filled circles show visible goal sessions. Dashed line indicates the 95th percentile of scores during baseline sessions. 18 of 168 post-injection sessions exceeded this criterion. (C) Thigmotaxis as a function of theta score. Animals could show normal levels of thigmotaxis during substantial reductions in theta power (data points in the left part of the graph, under the dashed line). (D) As in (A) for maximum (left) and minimum (right) mobility score. (E-F) As in B-C for mobility score. Dashed line is 5th percentile of mobility scores during baseline sessions. 7 of 168 sessions were below this criterion. In total, 23 of 168 sessions were excluded due to mobility or thigmotaxis criteria.

In the hidden goal version of the task, navigation accuracy was severely impaired after injection of either tetracaine, muscimol, or gabazine (Figure 7B). These deficits were due to inability to navigate based on spatial information rather than to general effects on locomotion or behavior (Figure 7D-F, Figure 8). Tetracaine and gabazine produced their strongest effects immediately after injection, and performance recovered within one hour. The rapid recovery demonstrates that animals retained knowledge of the target location, but were unable to retrieve and exploit that information to guide behavior. By contrast, place accuracy was impaired over a longer time period following muscimol injections, consistent with the longer period of theta power reduction produced by muscimol. Thus, regardless of injection type, deficits in locating the hidden goal were greatest when theta oscillations were most attenuated (Figure 7C).

To assess whether place accuracy deficits could be explained by locomotor or motivational impairments, we tested the same animals under identical conditions except the target location was marked with a visible cue. In the visible goal version of the task, behavioral performance was unaffected in any post-injection session, and there was no correlation between hippocampal theta power and task performance (Figure 7D-F). The fact that the animals could perform normally after disruption of hippocampal theta oscillations in a condition that does not require navigation based on allocentric cues eliminates the possibility that the behavioral deficits found in the hidden goal version of the task were due to non-specific drug effects. The profound impairment in spatial navigation after gabazine injection stands in stark contrast to its lack of effect on the place cell representation and indicates that the theta rhythm is critical for the behavioral readout of spatial information encoded in place cell activity.

## Discussion

### Differential effects of tetracaine, muscimol and gabazine on local field potentials

To dissociate the overall activity of hippocampal neurons from their rhythmic modulation, we selected three pharmacological agents that interfered with hippocampal rhythmic activity when injected into the medial septum, but which acted through different mechanisms. Local anesthetics (Mizumori et al., 1989; Koenig et al., 2011) were previously shown to affect the discharge of hippocampal place cells; comparisons with our work are made below. Lidocaine and the GABA-A receptor agonist muscimol reduce or eliminate the triangular firing field array of medial entorhinal grid cells (Brandon et al., 2011; Koenig et al., 2011). The third drug, the GABA-A receptor antagonist gabazine, was chosen based on previous work showing that GABA-A receptor blockade reduces rhythmic firing in medial septum (Brazhnik and Fox, 1999). As demonstrated here, gabazine reliably suppresses theta for intervals sufficiently long to properly sample place cell discharge.

### Cholinergically mediated hippocampal effects of septal GABA-A blockade

We observed reduced firing rates and degraded spatial coding after septal inactivation but not septal disinhibition, despite equivalent or greater reduction in hippocampal theta. Our results therefore indicate that factors other than rhythmic modulation of septohippocampal input determine place cell firing. Specifically, the medial septum is the primary source of cholinergic innervation to the hippocampus (Lewis and Shute, 1967; Frotscher and Leranth, 1985), which is effectively eliminated upon septal inactivation. Acetylcholine (ACh) release is closely associated with the theta state and place cell activity: the firing rates of cholinergic septohippocampal neurons (and hippocampal ACh levels) increase during theta (Dudar et al., 1979; Bianchi et al., 2003; Zhang et al., 2010) and ACh is necessary for normal place field activity (Ikonen et al., 2002; Brazhnik et al., 2003). By antagonizing GABA-A receptors, picrotoxin increases the activity of most septal neurons (Brazhnik, 2004), in line with the finding that septal bicuculline increases hippocampal ACh release (Moor et al., 1998; Chang et al., 2006; Roland and Savage, 2009). GABA-A blockade also reduces the rhythmicity of septal long spike cells (putative cholinergic septohippocampal neurons (Brazhnik and Fox, 1999). Thus, under GABA-A blockade the hippocampal theta oscillations are greatly diminished while delivery of ACh is enhanced. This pattern contrasts strongly with the effects of the other two drugs that reduce septal activity, each by a different mechanism. Our finding thus reveals novel differences between the selective reduction of the oscillatory component of the hippocampal theta state compared to elimination of both theta oscillation and the theta-associated increase in ACh delivery.

Although septal gabazine potently interferes with theta oscillations, its effects on other hippocampal oscillations differ from those of tetracaine and muscimol. Fast gamma oscillations were preserved or even strengthened following gabazine injections, in contrast with the broad decrease in gamma power following septal inactivation. Since ACh agonists promote gamma oscillations *in vitro* (Fisahn et al., 1998), these observations independently support the idea that hippocampal ACh release is elevated after gabazine injection. The preservation of fast gamma oscillations after gabazine infusions in the medial septum could allow the persistence of strong synchrony between the entorhinal cortex and hippocampus to support high-precision location-specific place cell firing. The concurrent reduction of slow gamma is consistent with the notion that it is mediated by interactions with CA3 (Colgin et al., 2009) via glutamatergic transmission that is suppressed by ACh (Hasselmo and Schnell, 1994). It seems likely that the preservation of the location-specific firing of place cells after gabazine also results from a preserved or enhanced septohippocampal release of ACh, since previous work has shown that muscarinic blockade degrades place cell firing patterns (Brazhnik et al., 2003; Brazhnik et al., 2004).

### Differential effects of tetracaine, muscimol and gabazine on place cells

The reduction of theta frequency modulation in place cell autocorrelograms following the infusions of each of the three drugs is approximately proportional to the drugs’ effects on theta amplitude. Strikingly, however, gabazine induced the smallest changes in the location-specific discharge of hippocampal place cells according to overall firing rate, spatial information, coherence and stability (measured by between-session similarity of rate maps). This dissociation of theta power and modulation by location implies that theta oscillations play at most a small role in processes that activate the representation of a familiar environment, in the ability of individual place cells to maintain their firing fields, or in the updating of activity based on translocation. Residual theta oscillations in some cells may support their location-specific firing, but no rhythmic activity was seen in many cells whose spatial firing properties were hardly changed by gabazine injections. Overall, our data indicate that the spatial and rhythmic firing patterns of place cells are largely independent properties.

The effects of medial septal injections of local anesthetics and muscimol on place cells were investigated twice before, with different outcomes. Using tetracaine, Mizumori et al (1989) reported large reductions in the firing rate of CA3 but not CA1 place cells. With a different local anesthetic, lidocaine, Koenig et al (2011) saw consistent decreases of place cell rates in CA1 and, with a small sample, little change in CA3. We found that both tetracaine and muscimol reduced place cell firing rates and disrupted spatial firing patterns in CA3 and CA1, with a somewhat larger decrease in CA3. The origin of the variable effects is unclear, but it could be due to differences in cannula placement, which was most dorsal in Mizumori et al (1989), most ventral in our study, and in between in Koenig et al (2011).

### Theta blockade and spatial problem solving

We asked how septal injections of tetracaine, muscimol and gabazine affect performance in two versions of a navigational task; in one version, the goal is visible, in the other it must be found by self-localization relative to stable landmarks. Performance in the visible version was intact regardless of which drug was injected. Thus, theta blockade did not interfere with memory for the basic rules of the task including the need to approach the goal; with locomotion; or with motivation. In contrast, each drug greatly impaired performance in the hidden goal version over a time course parallel to the drug’s effect on theta power, and behavioral performance strongly correlated with the power of the theta rhythm across sessions irrespective of the drug or mechanism.

Behavioral deficits following septal inactivation may be largely explained by degraded place cell activity and weak self-localization signals. However, the dissociation between the well-preserved location specific firing of hippocampal place cells and the navigational deficit induced by gabazine indicates that place cell activity alone cannot support accurate navigation. It is as if the absence of theta uncouples spatial information from the computations needed to select paths to desired locations. Though we focused primarily on drug effects on hippocampal theta power, other effects may contribute to the observed behavioral deficits. First, since all three drugs suppress both theta and low frequency gamma, the utilization of place cell output may depend on a gamma frequency synchrony between the CA3 and CA1 place cells (Colgin et al., 2009). Second, if septal gabazine, along with lidocaine (Koenig et al., 2011) and muscimol (Brandon et al., 2011) eliminates the triangularly arranged firing fields of medial entorhinal grid cells, the behavioral deficit might be ascribed to extra-hippocampal abnormalities. Indeed, since overall grid cell activity is also affected by septal inactivation, septal gabazine injections could be used in future studies to more directly test the role of theta rhythm in producing grid cell spatial firing patterns. However, recent work has demonstrated that spatial information in hippocampal output structures explicitly depends on phase-coding relative to hippocampal theta (Tingley and Buzsaki, 2018). Thus, the coherent theta rhythm, coordinating activity across hippocampus and associated downstream regions, may form a channel for the broadcast of spatial information that is critically necessary to convert spatial codes to adaptive behavior.

## Methods

### Subjects

Subjects were 11 adult male Long-Evans rats (Taconic) weighing 300-400 grams. Animals were singly housed, had free access to water, and were on a 12-hour light/dark cycle.

### Apparatus

Most recordings were made in a gray cylinder 76 cm in diameter and 50 cm tall. Two cue cards, one black and one white, were on the cylinder wall to provide orienting cues. Each card occupied 45 degrees with their closest edges 90 degrees apart. The floor was covered with gray backdrop paper that was replaced before each session. The arena was surrounded by a black curtain and lit by four small bulbs. The arena was inside a sound-attenuated box.

### Foraging behavior

Before surgery, rats were handled and food deprived to 85% of ad lib weight. They were then placed in the arena and trained to forage for 25 mg sugar pellets dropped from above to random locations at a rate of ~2/min. Training was done once per day for 1 week, during which rats learned to explore the whole cylinder, eating all pellets.

### Microelectrode and microinjection assembly

Recording implants held 8 individually drivable tetrodes. Each tetrode was wound with four 25 µm Formvar-insulated Nichrome wires (California Fine Wire, CA). The implant also had a 22 Ga stainless steel guide tube for medial septal injections. A 30 Ga stainless steel dummy cannula extended 1.5-mm past the end of the guide tube. Prior to use, tetrode tips were cut and gold-plated so the impedance of each wire was 30-80 kΩ.

### Surgery

After Nembutal anesthesia (45 mg/kg), the scalp was shaved and the rat placed in a Kopf stereotaxic frame. A midline incision was made and the skull leveled. Eight screws were put around the skull edge, two above the cerebellum. A 2 mm midline hole was made 0.7 mm anterior to bregma, over the medial septum. Two 2 mm diameter holes were drilled over the right hippocampus, one at −3.5 mm AP and 3.4 mm ML, the other at −4.3 mm AP and 4.1 mm ML away from bregma.

After removing the dura, the sagittal sinus above the medial septum was pulled to one side as the implant was lowered so that the guide cannula reached 5.0 mm below bregma; the dummy cannula was 6.5 mm below bregma. Tetrodes were individually adjusted to follow the curve of the hippocampal fields such that four tetrodes were 0.4 mm above the CA1 pyramidal layer and four tetrodes were 0.4 mm above the CA3 pyramidal layer. Ground wires were soldered to the cerebellar screws and the implant attached to the skull with dental cement. The wound was cleaned with 3% hydrogen peroxide and iodine and topical antibiotic was applied. Animals were allowed one week to recover from surgery with *ad libitum* food and water before screening began.

### Drugs and drug injection

Drugs were dissolved in sterile phosphate buffered saline (PBS; Fisher Scientific, BP2438). 4% tetracaine solution (151.31 mM) was prepared by dissolving 0.04 g (Sigma T3812) in 152 μl 1N HCl to which was added 848 μl sterile PBS. Muscimol (Sigma M1523) and gabazine (Tocris SR95531 HBr) were dissolved in sterile PBS to produce 1 μg/μl stock solutions and then diluted to the desired concentration. The final concentration of muscimol was 0.5 μg/μl or 4.38 mM. The final concentration of gabazine was 62.5 ng/μl or 168 μM.

Rats moved freely in their home cage during injections. Drugs were infused with an injection pump fitted with a 2.0 µl Hamilton syringe. Tygon tubing (~80 cm) was filled with PBS and attached to the syringe; a 30 Ga stainless steel injection cannula was on the other end of the tubing. The syringe plunger was retracted to draw 0.5 μl of air into the tubing followed by 1.5 μl of drug solution. The air bubble ends were marked. Injections took 2 min at 0.25 μl/min for a total volume of 0.5 μl. The injection cannula was left in place for 3 min more. The injection was successful if the air bubble shifted by the distance corresponding to 0.5 μl. If this criterion was not met, the injector was gently raised and lowered within the guide tube and a second attempt was made. If the second attempt was not successful, the experiment was postponed. After injection, the dummy cannula was replaced and the rat returned to the cylinder.

### Electrophysiology; screening and data acquisition

After recovery from surgery, rats were screened for action potentials and local field potentials during foraging. Single unit data were filtered from 300 to 5000 Hz. If the amplitude on any wire of a tetrode exceeded a threshold, 1.0 msec of data sampled at 33 μs intervals from all 4 wires were time-stamped and stored for off-line analysis (Cheetah System; Neuralynx Inc., Bozeman, MT). For local field potentials (LFP), the signal was band-pass filtered from 1 to 250 Hz and sampled at 3030.30 Hz. Tetrode tips were advanced 60 microns/day and each channel monitored for complex-spikes, indicating entry into the CA1 or CA3 pyramidal layer. To assist tetrode positioning, each channel was also monitored for LFPs referred to the cerebellar screws. Entry into the CA1 pyramidal layer was indicated by the presence of complex-spike cells, reversed sharp-wave polarity and the beginning of a theta phase shift.

A single tetrode was driven deep into stratum radiatum and used to record the LFP. Once this tetrode passed through the pyramidal layer, theta amplitude grew rapidly and phase shifted ~180°. The wire with the largest theta activity in a pre-injection session was used for further analysis.

### Experimental protocol: Single cell sessions

After a 16-minute baseline session, rats received 1 of 4 intraseptal injections: vehicle (sterile PBS), 4% tetracaine, muscimol (250 ng) or gabazine (31 ng). The first post-injection session started 7 minutes after the start of the injection and lasted 32 minutes. Three more 16-minute recording sessions were done at 60, 120 and 240 minutes post-injection.

Twenty-four hours after a successful injection, a recovery recording session was performed. Single unit and LFP signals were monitored for complex spikes and theta. If these were present, a new experiment was started. Each rat received at most 8 injections (two for each solution); often, fewer injections were done because single cell waveforms could no longer be detected during a recovery session. The order of injections for each animal is presented in Table 1.

**Table 1.** Injection sequence for each animal. Drugs are vehicle (V), tetracaine (T), muscimol (M), and gabazine (G).

| Animal ID | Injections |
| --- | --- |
| 923 | T |
| 925 | T |
| 949 | T |
| 954 | V, G, M, T |
| 955 | V, G, M, V |
| 957 | M, T, G, V, V, G, M |
| 961 | V, M, T, V, G |
| 962 | G, V, M, T, G, V, M, T |

### Perfusion and Histology

Once tetrodes passed through the pyramidal cell layer, the rat was deeply anesthetized with urethane. Current (20 µA for 20 sec) was passed through one wire of each tetrode to mark its location. Rats were perfused transcardially with 0.9% saline followed by 4% paraformaldehyde. Fixed brains were embedded in gel matrix, cut in 30µm sections on a cryostat, and stained with cresyl violet. We analyzed only data from rats in which the cannula was in the medial septum and the sets of 4 recording tetrodes were in CA3 and CA1.

### Running speed

The rat’s head position was tracked with an overhead TV camera by detecting lights on the headstage. A 24×24 grid of 3 cm square pixels were used to analyze spatial firing. Position was measured at 30 Hz. Running speed was calculated for each sample by finding the distance traveled in a 366 msec interval (5 samples on either side of the target sample). To characterize speed dependency of LFPs and single cell activity, speeds were averaged across 3 seconds.

### Local field potential analysis

Frequency spectrum analyses were performed in MATLAB (Mathworks, MA). For spectrogram visualization and power calculations, data were down-sampled to 757.58 Hz and the short-time Fourier transform was calculated with 1024-sample Hann windows with no overlap, zero-padded to 4096 samples. Power spectra were averaged in a sliding 16-minute window and *theta scores* were calculated within each window to determine the period of minimal theta to define the ‘post-injection’ analysis period. A line was drawn between the points at 5 and 12 Hz in the log-power spectrum and theta power was taken as the integral of the spectrum above the line. Theta scores were computed by dividing post-injection theta power values by the overall baseline theta power. Power in other frequency ranges was not typically confined to clear peaks. Nevertheless, to isolate specific oscillatory power from 1/f spectral noise, we fit a line to the baseline log-power spectrum between the points at 1 and 100 Hz and subtracted this line from the baseline and post-injection spectra.

### Analysis of single-cell data

Waveforms from candidate cells were clustered according to spike amplitude and principal components with Plexon Offline Sorter (Plexon Inc., Dallas, TX). To generate firing rate maps and place cell statistics, clusters were analyzed using custom MATLAB code (Mathworks, CA). In all analyses, data were selected for periods in which the rat moved > 3 cm/s. For each cluster, the number of spikes fired in each pixel was divided by the time spent in that pixel to calculate firing rate. Increasing firing rates in a pixel are coded in the color order: yellow, orange, red, green, blue, purple. White coded pixels unvisited by the rat.

Overall rate, *spatial information, spatial coherence* and *similarity score* were calculated 184 for each unit. Overall rate was computed as the number of spikes in a cluster divided by 185 recording duration. Spatial information (I_pos_) was calculated as in Olypher et al. 2003 using 186 ms time bins for computing spike count (*k*) distributions in each pixel (*x_i_*).

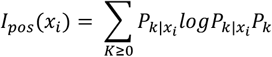

I_pos_ was averaged over all arena pixels for each cell to compute the mean spatial information, equivalent to mutual information between spike counts and position. Coherence is a measure of the smoothness of place cell spatial firing and was computed as the z-transform of the correlation between the rate in each pixel and its eight nearest neighbors. Discriminated waveforms were considered place cells if overall rate was between 0.1 and 4 Hz, coherence > 0.3 and the cell fired in < 50% of the pixels in the arena. Units with overall rate > 5 Hz were considered to be interneurons. Units not matching either of these criteria were excluded from analysis. Similarity scores compare the firing rate distribution in 2 sessions. Rate maps were constructed for each session; the similarity score was the z-transformed pixel-by-pixel correlation between these maps.

Autocorrelations for each cluster were computed and displayed to visualize rhythmic firing. 5-ms binned autocorrelograms were computed for ± 500 ms lags. The central peak was truncated to the highest value occurring at >50 ms lag, and the mean value of the autocorrelogram was subtracted. The power spectrum of the autocorrelogram was computed using a fast Fourier transform zero-padded to 2^16^ samples. The *findpeaks* function (MATLAB Signal Processing toolbox) was used to find a single peak between 6 and 11 Hz in the power spectrum. The *theta-modulation score* was computed as the ratio between the mean power within 1 Hz of the theta peak and the total power between 0 and 50 Hz (Yartsev et al., 2011).

### Place accuracy task – Training

The foraging behavior described above allows the assessment of the spatially selective activity of the hippocampal neurons since the animal uniformly visits the entire floor surface of the recording arena. To assess spatial memory and navigational ability we separately trained animals to navigate to a circumscribed area of the arena for food reward. We then evaluated the effects of tetracaine, muscimol, and gabazine on the rats’ ability to use spatial memory. The specificity of these effects was assessed by comparing the performance in the spatial task to performance in a task run in the same environment but which only required remembering cue-response associations.

Five animals were trained and tested in the place accuracy task. The rat’s position was tracked as usual and a sugar pellet released when a set of location criteria were met. To trigger release, the rat’s head had to remain inside an experimenter selected 16-cm diameter circle for 1.2 seconds. The rat then had to stay outside the target circle for 3 seconds before another reward could be released. During initial training the center of the target area was marked with an 8 cm diameter, 1.4 cm high disk. Two 20 min training sessions were run per day until each rat triggered > 40 releases in a session for 3 consecutive sessions. Next, sessions with no disk were alternated with disk sessions until the rat released at least 40 pellets for 3 consecutive no-disk sessions. Finally, only no-disk sessions were performed. When rats released at least 40 pellets for 3 consecutive no-disk sessions they were ready for injections. We refer to disk and no-disk conditions as visible and hidden goal variants of the task. Successful performance during the visible goal variant can be based either on following the disk or on spatial navigation to the disk; however, performance during the hidden goal task requires spatial navigation.

### Place accuracy task – Testing

Drug concentrations and the injection protocol were identical to those used in the place cell/LFP work. A rat was tested with each substance in the hidden goal version of the task. It was then re-tested with each substance in the visible version of the task. Local field potentials confirmed that substances affected theta as expected. Theta scores were computed as above, but from average power spectra over full 20-minute sessions. Behavioral sessions were excluded from analysis if animals showed excessive thigmotaxis or immobility. Thigmotaxis, the tendency to remain near the walls and away from the center of the arena, was measured as the percent time over the 20-minute behavioral session spent in the outer 10% of the arena diameter, and immobility score was the percentage of time where running speed was less than 5 cm/s.

An experiment consisted of four sessions identical to those during training. On the experimental day, a 20-minute baseline session was performed, followed by drug injection. A second 20-minute session was begun 7 minutes after the start of injection. The third and fourth sessions took place one hour and twenty-four hours after injection. Twenty-four additional hours were allowed to pass before starting a new experiment. Dwell time maps were constructed at 2 pixel/cm resolution and performance was measured as percent time in target area.

### Statistics

Statistical analyses were performed using MATLAB. Differences between drug conditions were detected with one-way ANOVA, or twoway ANOVA if the distinction between CA3 and CA1 was a factor. Post-hoc comparisons were made using unpaired t-tests. Within-subjects effects were detected with one-way repeated measures ANOVAs and post-hoc comparisons were made with paired t-tests. Multiple comparisons were corrected using the false discovery rate where appropriate.

## Acknowledgments

We thank B. Rivard and H. Tassinari for technical assistance; J. Barry, A. Fenton, J. Kubie and E. Song for helpful discussions and assistance with experimental setup; and C. Hull and K. Franks for reading early versions of the manuscript. Funding: This work was supported by funding from NINDS (NS20686) and NIMH (R21MH094946 and R21MH115421).

